# Drone-Based Monitoring of Reproductive Potential in a Foundational Shrub Species

**DOI:** 10.1101/2025.06.27.662010

**Authors:** Ryan Scott Wickersham, Megan E. Cattau, Jennifer S. Forbey, Valorie Marie, Andrii Zaiats, Donna M Delparte, T. Trevor Caughlin

**Affiliations:** Boise State University, Boise, ID, USA; Idaho State University, Pocatello, ID, USA

**Keywords:** Drones, sagebrush steppe, phenology, structure-from-motion, intraspecific variation, seed sourcing

## Abstract

Restoration and conservation of native plant populations will benefit from identifying individual plants with high reproductive success. While high-fecundity plants are ideal for seed sourcing, locating these plants across heterogeneous landscapes presents a logistical challenge. This challenge is especially significant for big sagebrush (*Artemisia tridentata*), a foundational species that is the focus of large-scale seed collection for restoration efforts in western rangelands. We evaluated whether cost-effective RGB imagery from unoccupied aerial vehicles (UAVs) could map flower stalk production in big sagebrush plants. Models were trained using three years of data from four sites spanning an elevational gradient that included all three big sagebrush subspecies: *A. t. wyomingensis, A. t. vaseyana*, and *A. t. tridentata*. Our model predicted flower stalk production from UAV imagery with a Mean Absolute Error (MAE) of ∼100 stalks, which is relatively low given that some plants produced more than 700 stalks. A hurdle model that explicitly accounted for excess zeroes outperformed simpler negative binomial models, suggesting that reproductive failure is distinct from flower stalk production in reproductive plants. Structural metrics, including height differences between June and September, canopy height, and edge-to-area ratio of plant crowns, had stronger effects in our model for counts of flower stalk production than spectral data. Model performance was consistent across environmentally heterogeneous sites but declined when applied to years excluded from training, indicating that year-specific training data may be necessary for interannual predictions. These results demonstrate that UAVs can monitor reproductive potential in wild plants and help identify high-fecundity individuals for seed collection. Our work underscores the need for future research that can improve predictions of flower production, including integrating multispectral data and increasing model reliability across years to support climate-resilient restoration strategies.

## Introduction

Restoring native plant populations depends on the ability to collect high-quality seed from wild plants (Pedrini et al. 2020). Seed collection can be challenging, especially in rangelands, which are vast, remote, and often topographically and environmentally complex (Roundy et al. 1997, Kildisheva et al. 2016). Native species in these systems also often exhibit strong local adaptation, making it risky to use seed from distant or genetically mismatched sources (Baughman et al. 2019). This challenge is particularly acute for species with wide distributions, where seed from the wrong source may lead to poor establishment, low survival, or long-term fitness declines (Shryock et al. 2017). In this context, identifying highly fecund individuals, plants that produce large quantities of viable seed, is a critical first step toward restoring resilient, locally adapted populations (Pedrini et al. 2020). These individuals can serve as key seed sources, but are often sparsely distributed and difficult to detect at the landscape scale.

Big sagebrush (*Artemisia tridentata*) exemplifies the challenges of seed collection from wild populations. As a foundational species of the western U.S., sagebrush plays a vital role in supporting biodiversity and ecosystem function but is increasingly imperiled by climate change (Dalgleish et al. 2011), more frequent and severe wildfires (Balch et al. 2017), invasive species (Smith et al. 2023) and human population growth (Requena-Mullor et al. 2023). These threats have prompted massive restoration efforts, including more than $100 million spent on sagebrush seed collection by the Bureau of Land Management since 1990 (Simler-Williamson and Germino 2022). Intraspecific variation between sagebrush plants is common and impacts both immediate restoration success and long-term prospects for population persistence in a changing climate (Richardson and Chaney 2018). Consequently, the difficulty of sourcing sagebrush seed from locally adapted populations can limit restoration success (Brabec et al. 2015).

Strategically targeting seed collection for highly fecund individuals will require understanding the demographic rates that drive seed availability, beginning with flower production. Flower production is a demographic rate that shapes plant fitness and population growth rates in unstable environments (Jacquemyn et al. 2010, Miller et al. 2012). High rates of flower production can indicate local adaptation in plant populations, resulting in greater overall fitness (Simón-Porcar et al. 2021). In big sagebrush, flower stalk production varies between subpopulations (Richardson et al. 2017), leading to gene by environment interactions in seed yield that influence how subpopulations will respond to climate change (Richardson and Chaney 2018). These previous studies by Richardson et al. used climate variables to interpolate results from common garden experiments across large spatial extents; directly measuring flower stalk production at scale would provide complementary data to test hypotheses for local adaptation and target seed collection to individual plants.

Advances in remote sensing technology offer a promising solution for assessing plant fecundity at scales far exceeding field measurements. Unoccupied aerial vehicles (UAVs), in particular, can collect imagery with relevance to plant structure and demography (Torresani et al. 2023, Olsoy et al. 2024). Precision agriculture has leveraged these new technologies to improve crop yields, including mapping flower production in agricultural fields (Wan et al. 2018, Yang et al. 2022), Similarly, high-resolution remotely sensed data has shown potential to predict flower production in wild plant populations, such as UAV-based predictions of flowering in tropical tree canopies (Lee et al. 2023). However, whether remote sensing techniques pioneered on agricultural crops or canopies of large trees can predict flower stalk production for rangeland plant species in natural landscapes remains an open question.

Sagebrush, in particular, presents several challenges for remote sensing applications. Differentiating the spectral signal of vertical, gray sagebrush leaves from other landscape features, including bare ground and biocrust, is not straightforward (Mitchell et al. 2012). One potential solution to this challenges is structure-for-motion (SfM) derived from UAV imagery. SfM involves capturing high-resolution RGB images with cameras flown at low altitudes with high forward and side overlap. These overlapping photos enable 3-D reconstruction of shrub canopies, producing dense point clouds that facilitate precise structural measurements (Olsoy et al. 2024). Leveraging SfM technology, UAV imagery has the potential to overcome mapping challenges in semi-arid ecosystems, enabling accurate detection of individual sagebrush canopies for remote sensing applications (Howell et al. 2020). The genetic diversity of sagebrush also complicates remote sensing studies. The big sagebrush species complex includes multiple ploidy levels and genetically distinct subspecies (Grossfurthner et al. 2023). These genetic differences not only result in separate ecological niches, but also contribute to spectral dissimilarity among sagebrush populations (Robb et al. 2022). Spectral differences among populations may limit the ability to develop generalizable models for predicting sagebrush characteristics, including flower stalk production.

In this study, we develop UAV-based models to estimate flower stalk production in big sagebrush across a landscape characterized by high intraspecific diversity in subspecies and ploidy level (Grossfurthner et al. 2023). Our approach leverages high-resolution RGB imagery and SfM point clouds to quantify canopy structure and spectral characteristics of individual shrubs. We addressed three key objectives. First, we developed a model to predict flower stalk production, including identifying the spectral and structural variables most strongly associated with reproductive output. Second, we assessed whether including UAV-derived data from multiple points in the growing season—specifically, canopy height differences between early and late season flights—improves model performance. Third, we tested the spatial and temporal transferability of our models by evaluating their performance when applied to new sites and years not included in model training. Models that generalize well across space and time are essential for operationalizing UAV-based monitoring tools across diverse landscapes and fluctuating environmental conditions (Blackburn et al. 2025). Together, these objectives advance the development of scalable, low-cost methods to map native plant fecundity in rangeland ecosystems.

## Materials and Methods

### Study Site

We conducted our research at Castle Rocks State Park located within the Northern Great Basin ecoregion (42.1367 N, 113.6769 W). Across our study landscape, elevation ranged from 1624-1840 meters. Castle Rocks has 25 different soil types with dominance by Andisols, Aridisols, and Oxisols (Keck et al. 2020). Precipitation averages at the site range from 251-460mm (Keck et al. 2020, Northern Great Basin Ecoregion Rapid Ecoregional Assessment Final Report. 2013). Subspecies of big sagebrush present in our study landscape include Wyoming Big Sagebrush (*Artemisia tridentata wyomingensis*), Mountain Big Sagebrush (*A. t. vaseyana*), and Basin Big Sagebrush (*A. t. tridentata*). Within our study landscape, we placed four 50x50 meter plots across an elevational gradient, stratified to represent different subspecies and ploidy levels (Fig. 1).

**Figure 1.**
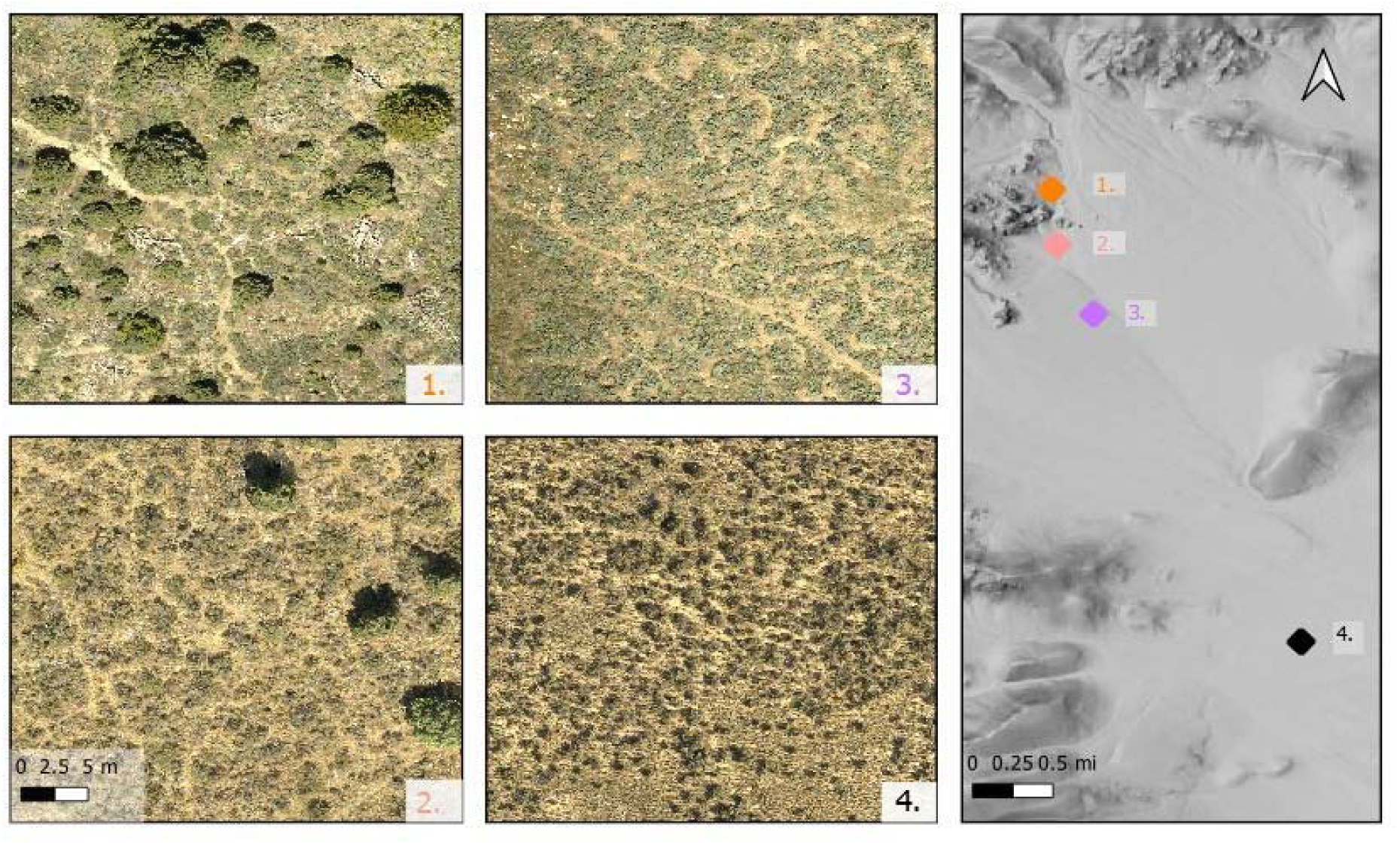
Site map at Castle Rocks State Park study location. From top to bottom are site 1 (1840m, dominated by *A. t. vaseyana*), site 2 (1780m, dominated by *A. t. vaseyana*), site 3 (1750m, dominated by *A. t. tridentata*), and site 4 (1620m, dominated by *A. t. wyomingensis*). Each site is 50x50 meters. Color panels represent RGB imagery collected by Unoccupied Aerial Vehicles (UAV). The map on the right panel represents landscape-level topography (ESRI).

### Demographic field measurements

Over the course of three years from 2021-2023, we collected field data for our four plots during peak flowering season in mid-September. Within each plot, we randomly tagged 50 big sagebrush plants for demographic measurements. We supplemented these core plants with measurements of flower stalk production at additional, opportunistically selected plants within the footprint of UAV imagery, for 1009 total measurements of flower stalk production. We measured flower stalk production by counting newly produced flower stalks, defined as inflorescences separated at the first stem node. We were able to distinguish current year flower stalks from previous years’ flower stalks, as newly produced stalks were green and pliable. After measurements were made, we recorded precise GPS coordinates of each shrub sampled using a survey-grade RTK GPS unit (Topcon HiPer V, Topcon Positioning Systems Inc., Livermore, CA, USA), with ∼0.02 m accuracy.

### Image collection and processing

We collected UAV imagery using a *DJI Mavic 2 Pro* with a 20MP Hasselblad RGB sensor. All flights were conducted by Section 107 FAA licensed pilots and in accordance with all state, federal, and local regulations. Flights were conducted between 1000-1400 (solar noon) in June and September of 2021-2023. We set flight parameters to produce pixels with a resolution of 1 cm^2^. We designed a flight plan with a cross-grid pattern and a single diagonal pass to create a highly detailed structure-from-motion (SfM) model for reconstruction of shrubs in the point cloud. Resulting imagery was processed to produce RGB orthomosaics and SfM point clouds using Open Drone Map, an open-source software platform (Marie et al. 2023). For additional details on flight protocol and processing parameters see Appendix S1.

### Canopy height model and segmentation

We used SfM point clouds to create a Canopy Height Model (CHM), using point cloud filtering software *CloudCompare*. *CloudCompare* applies a cloth simulation filter to discriminate between ground and non-ground points, enabling interpolation of the CHM. The resultant CHM consists of 1 cm^2^ pixels, each with a canopy height value. We then applied the lidR package (Roussel et al. 2024) in the R programming language to segment individual plant canopies in the CHM. We first used the lmf algorithm (Popescu and Wynne 2004) to identify points representing the top of big sagebrush canopies. We then used the silva2016 algorithm (Silva et al. 2016) to segment pixels in the neighborhood around canopy tops (Fig. 2). We have found that both of these algorithms perform well in identifying sagebrush canopies from SfM-derived canopy height models (Zaiats et al. 2024).

**Figure 2.**
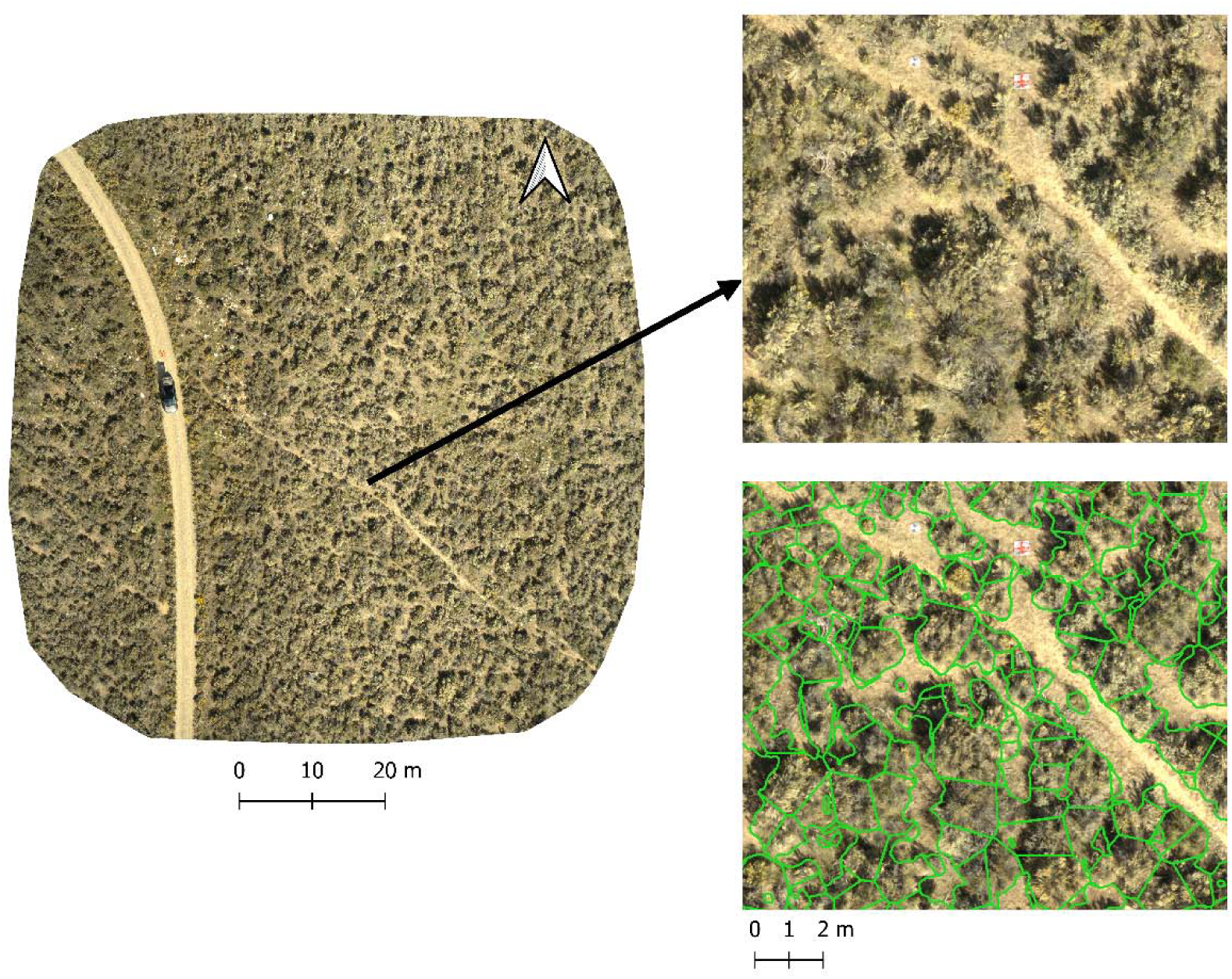
Segmentation results from site 3 at Castle Rocks State Park. The left image represents RGB drone imagery, acquired in 2023. The black arrow points to a zoomed in area of the drone image and the bottom right panel displays segmentation results, with green polygons representing segments.

### Combining segments with field data

The end goal of data processing was to create a table with a column for field-measured flower stalk counts, matched with columns for remotely sensed features of sagebrush canopies. To produce this data table, we merged segmented canopies with field data and their respective GPS points.

We created structural variables by aggregating canopy height model pixels for each sagebrush canopy. We expected larger plants to produce more flower stalks (Caughlin et al. 2019), and represented plant size in our models with mean canopy height within segments. To represent canopy heterogeneity, including heterogeneity created by flower stalks growing over canopies, we also included canopy height standard deviation. Structural metrics also included the 2-dimensional geometry of the shrub, represented as edge-to-area ratio, a metric frequently used in object-based classifications (Jochems et al. 2021).

We calculated canopy height and edge-to-area ratio from September data, the same month as field measurements of flower stalks. We also expected within-season growth to correlate with flower stalk production, as newly produced flower stalks add height to the canopy height model. To represent seasonal growth, we calculated pixel-level differences in total canopy height between the June and September SfM products for each year. We then aggregated difference pixels to sagebrush crowns, producing metrics for mean height difference and the standard deviation of height differences for each plant.

In addition to structural covariates, we also included RGB bands as predictors of flower stalk production. For these bands, we included the mean spectral reflectance for each shrub as well as the standard deviation within shrub canopies. Previous studies have demonstrated the importance of spectral bands for measuring flower production (Lee et al. 2023). Because the red and green bands were very highly correlated (Pearson’s *r* >0.98), we only included red and blue bands to develop model covariates.

Finally, we included covariates representing both year and location within one of our four study sites. Because our dataset spans three years that differed in precipitation and temperature, we included year as a categorical covariate in our models. We also included site identity as a categorical variable to account for spatial variation—such as differences in elevation and soil properties—that influences sagebrush demography (Dalgleish et al. 2011).

### Model development

We used generalized linear models (GLMs) in a Bayesian framework to predict flower stalk production, assuming a negative binomial distribution to account for overdispersion, which is common in ecological data (Bolker 2008). Given the high proportion of zeros—plants that did not produce any flower stalks in a given year—we also evaluated zero-inflated and hurdle models. The zero-inflated model assumes zeros arise from two sources: structural zeros, due to reproductive barriers (e.g., poor growing conditions or lack of pollination), and sampling zeros, due to stochastic variation. This model then combines both zero types in a single probability distribution. In contrast, hurdle models treat all zeros as arising from a single binary process, modeled separately from the count distribution for non-zero observations (Feng 2021). Consequently, hurdle models can incorporate covariates to explain both whether or not a plant reproduced, and if so, how many flower stalks the plant produced. Our first step in model development was to compare the predictive performance of negative binomial, zero-inflated negative binomial, and hurdle negative binomial models.

Model covariates included edge to area ratio, the mean and standard deviation of canopy height, red, and blue pixels within plant crowns, the year of data acquisition, and the mean difference in canopy height between June and September within a year. Because we expected that the effect of covariates, as well as baseline flower stalk production, would differ between sites with different environmental conditions and sagebrush genotypes, we also included site identity as a random intercept and all covariates as random slopes, conditional on site identity. For the hurdle negative binomial model, we used the same model structure for both the zero and non-zero components of the model.

To directly compare covariate effect sizes, we placed covariates on the same scale by centering covariates around the mean and dividing them by two standard deviations (Gelman 2008). We fit models using the **brms** package in R (Bürkner et al. 2024) and assessed model convergence by visually examining HMC output and by using model convergence metrics (R-hat and effective sample size) provided by brms. We evaluated model fit using a Bayesian version of the R^2^ metric and Mean Absolute Error (MAE), calculated out-of-sample using leave-one-out cross-validation.

### Evaluating model predictive capacity

A main goal of our study was to assess which types of data are most important for predicting flower stalk production. Because conducting UAV flights twice in a single year increases the cost of data collection, we compared predictive accuracy for our full model, including variables representing canopy height differences between June and September, to a model with data from September only. Because collecting field validation data is expensive and time-consuming, we also aimed to assess how well our model performs when predicting flower stalk production using a model trained on a different year. If models perform well, regardless of whether training data included year-specific field data, results would suggest that minimal field data is required to apply our models to new imagery. Poor performance for out-of-year sampling would indicate that some amount of new field validation data is necessary when applying our model to different years. We also evaluated the model’s ability to generalize across space by implementing a leave-one-site-out cross-validation approach. In this spatially explicit test, we trained the model using data from all but one site and evaluated its performance on the held-out site, rotating through each site in turn.

## Results

### Data summary

After matching segmented crowns to field GPS points, we had a total sample size of 561 plants. Across all years and sites, we measured a mean (± SD) of 76 (±124) flower stalks per plant, with a range from 0-709 flower stalks per plant. Across all years and sites, the proportion of plants with zero flower stalks was 20.32%. Variation in flower stalk production was greater across sites than years (Fig. 3), with plants at the lowest elevation site producing nearly three times as many flower stalks, on average, compared to the highest elevation site. However, variation in the proportion of zeroes was greater between years than sites, including nearly a 30% difference in plants with reproductive failure between 2021 (37.7% of plants) and 2023 (9.8% of plants).

**Figure 3.**
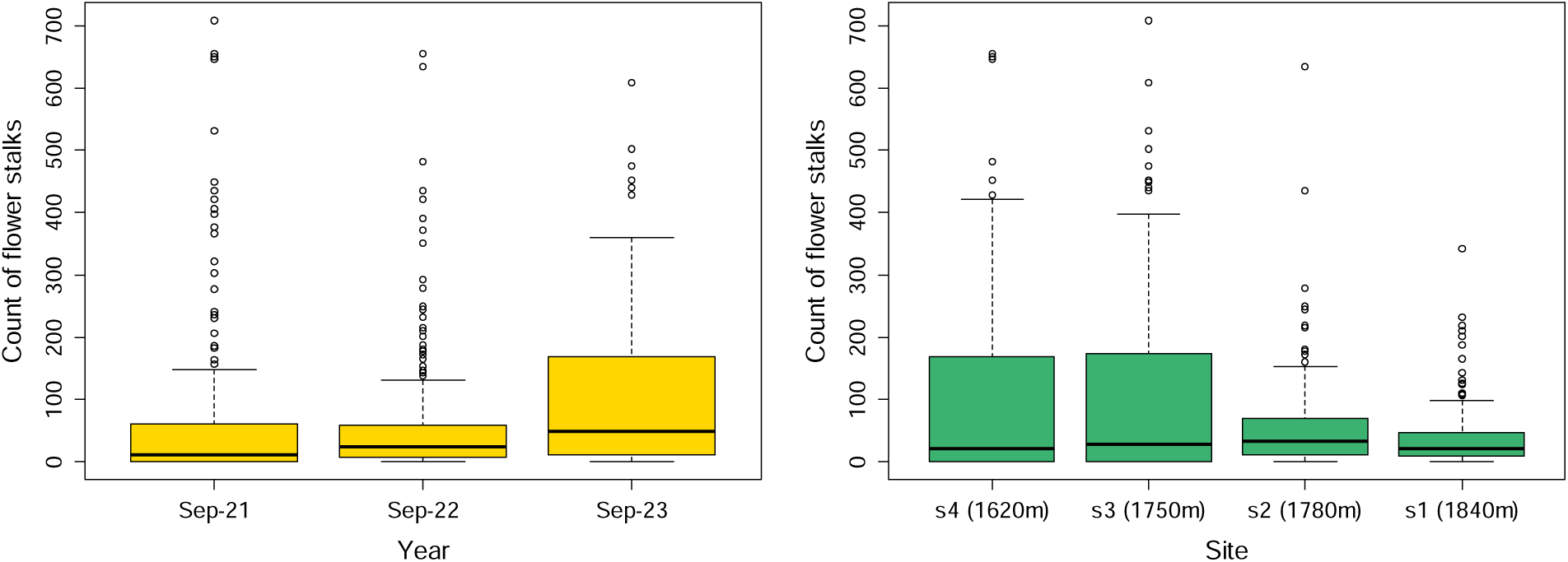
Data on sagebrush flower stalk production, aggregated by year (left panel) and site (right panel). Boxes represent the interquartile range (from the first to the third quartile), with the thick black line indicating the median. Whiskers extend to the smallest and largest values within 1.5 times the interquartile range from the lower and upper quartiles, respectively. Dots are outliers.

### Predictive ability of negative binomial models

The full model including data from all years, all sites, and seasonal differences in canopy height, had a Bayesian R^2^ of 43% (90% CI: 33–50%). Out-of-sample MAE (mean absolute error) from four-fold cross-validation, revealed a median error of 100 flower stalks (90% CI: 89 –126 flower stalks). In contrast, the standard negative binomial model (without zero inflation) performed substantially worse, with a median MAE of 136.6 and much more uncertainty (90% interval: 107–421 flower stalks). The zero-inflated negative binomial model showed intermediate performance (median MAE = 114.2; 90% CI: 97–196 flower stalks), but was still less accurate than the hurdle negative binomial model.

### Effects of covariates in full model

In the count (non-zero) component of the hurdle model—representing how many flower stalks were produced by plants that successfully reproduced—structural variables were the most influential predictors (Fig. 4). The strongest effect was the edge-to-area ratio of plant crowns, which had a relatively large negative estimate (-1.0 with 90% CI from -3.62 to 1.69), though the wide credible interval indicates substantial uncertainty. This result suggests that more fragmented or irregular plant crowns may be associated with reduced reproductive output. Two other structural variables—maximum difference in canopy height (Max.difference) and mean canopy height (Mean.CHM)—showed positive effects on flower stalk production, supporting the idea that more structurally variable and taller vegetation is associated with greater reproductive effort.

**Figure 4.**
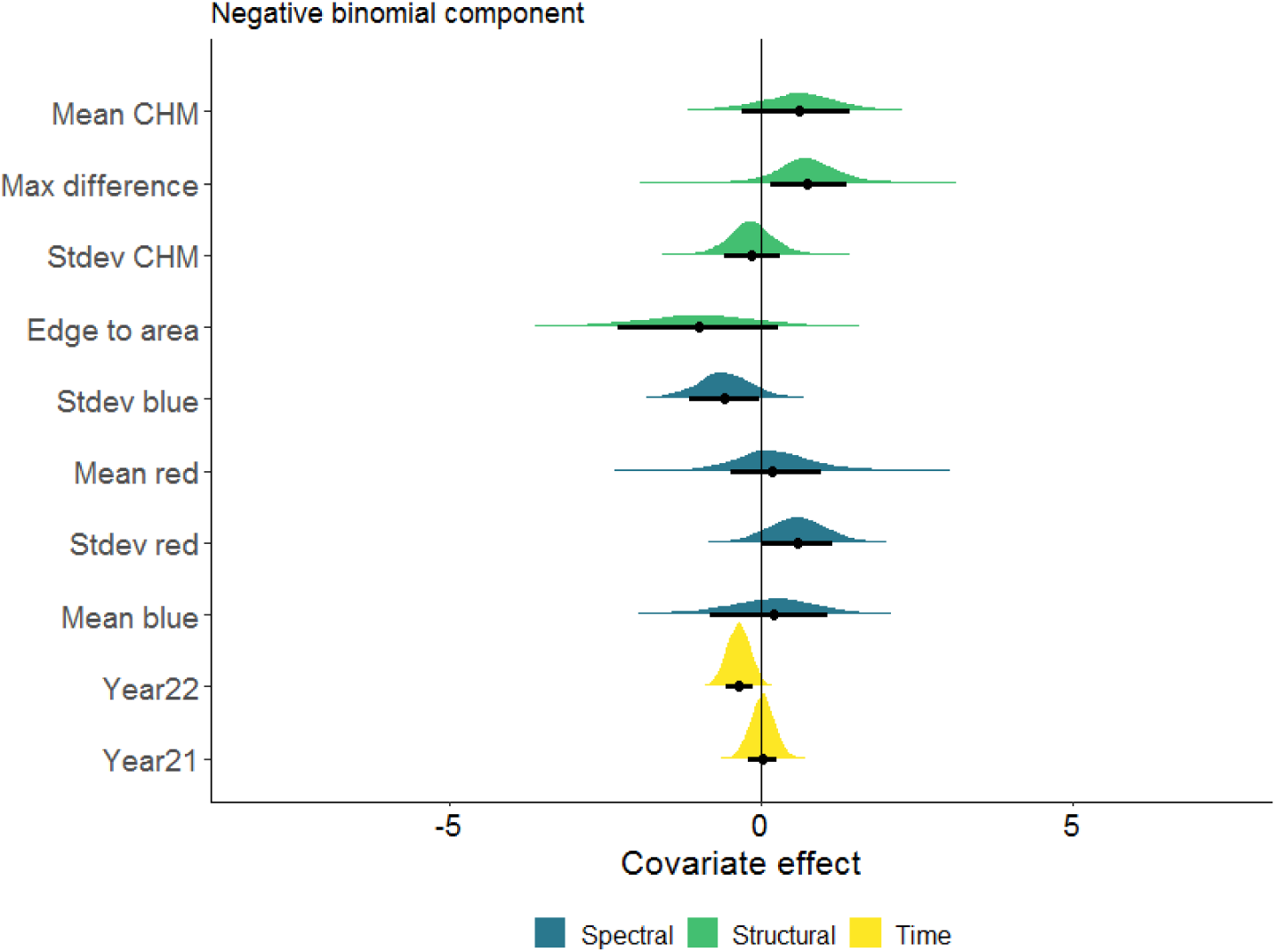
Effect of covariates on the count of flower stalks. This figure displays output from the negative binomial component of the hurdle model, representing only plants that successfully reproduced. Each hump represents posterior draws for a given covariate. Black dots represent median estimate and black lines represent 80% credibility intervals.

Among the spectral predictors, variables representing texture (i.e., the standard deviation of red and blue pixel values: std_red, std_blue) had larger absolute effects than those representing average color (e.g., Mean.red, Mean.blue). This pattern implies that spectral heterogeneity within plant crowns may be more informative than simple spectral means for explaining reproductive output.

For the zero component of the hurdle model, representing whether or not a plant produced flower stalks, the yearly effect for 2021 was the strongest predictor variable (Fig. 5). Aside from yearly effects, both spectral and structural components were important predictors of reproductive probability. The most important spectral variable for predicting whether or not a plant reproduced was the mean of blue pixels within plant crowns, while the most important structural variable was the standard deviation of canopy height pixels within plant crowns. In general, variables that had a strong effect on the count of flower stalks in the model’s non-zero component had weak and uncertain effects on whether or not a plant reproduced. An exception was the effect of mean canopy height, which was positive for the model’s non-zero component and negative for the zero component. This indicates that, on average, taller plants were less likely to reproduce but those taller plants that did reproduce tended to have more flower stalks.

**Figure 5.**
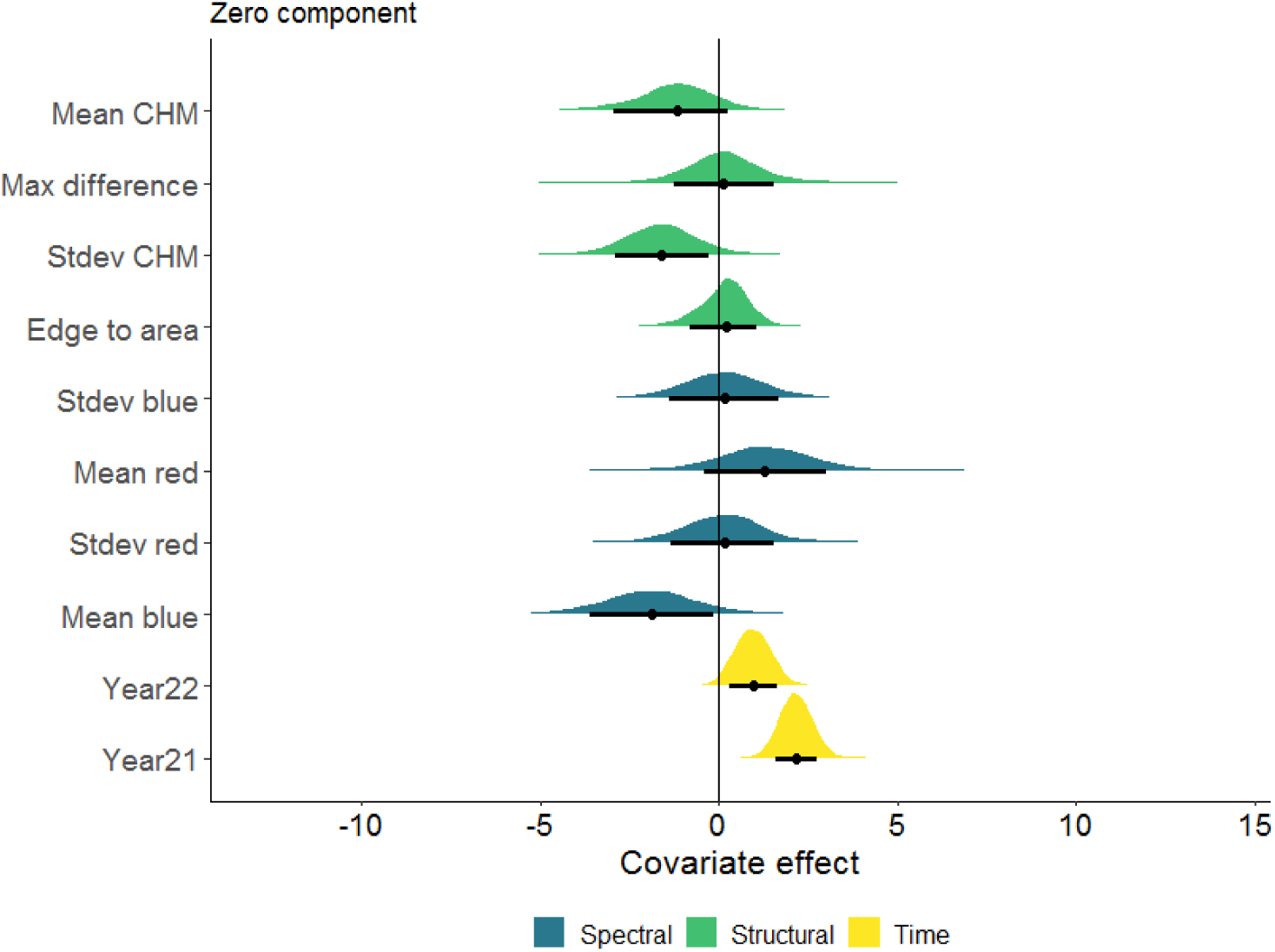
Effect of covariates on the probability of reproduction. This figure displays output from the zero component of the hurdle model, representing whether or not plants successfully reproduced. Each hump represents posterior draws for a given covariate. Black dots represent median estimate and black lines represent 80% credibility intervals.

### Predictive importance of seasonal height differences

We compared models with and without seasonal differences in mean canopy height to assess whether an additional UAV flight in June improved predictions of flower stalk production. The model without seasonal canopy difference variables yielded a mean absolute error (MAE) of 96 flower stalks (90% CI: 88 to 108), which was slightly lower than the MAE of 100 flower stalks (90% CI: 89 to 126) for the model that included seasonal canopy height differences. This indicates marginally better out-of-sample performance for the reduced model. Predictive uncertainty was also lower in the reduced model, as reflected in a narrower credibility interval. These results are borne out by visual examination of predicted vs. observed values for both models (Fig. 6). Both models tended to underpredict flower stalk counts, with an average bias of -30 flower stalks for full model, compared to an average bias of -32 flower stalks for the model without canopy height differences). Both models also included one prediction for high flower stalk production for a plant that produced zero flower stalks in reality. Despite the slightly better out-of-sample performance of the simplified model, the Bayesian R² was higher for the full model that included canopy height differences, with a median of 43% (90% CI: 36% to 49%) compared to 38% (90% CI: 30% to 45%) for the reduced model. This result suggests that the additional spring flight may have captured useful explanatory information, even if it did not improve predictive accuracy.

**Figure 6.**
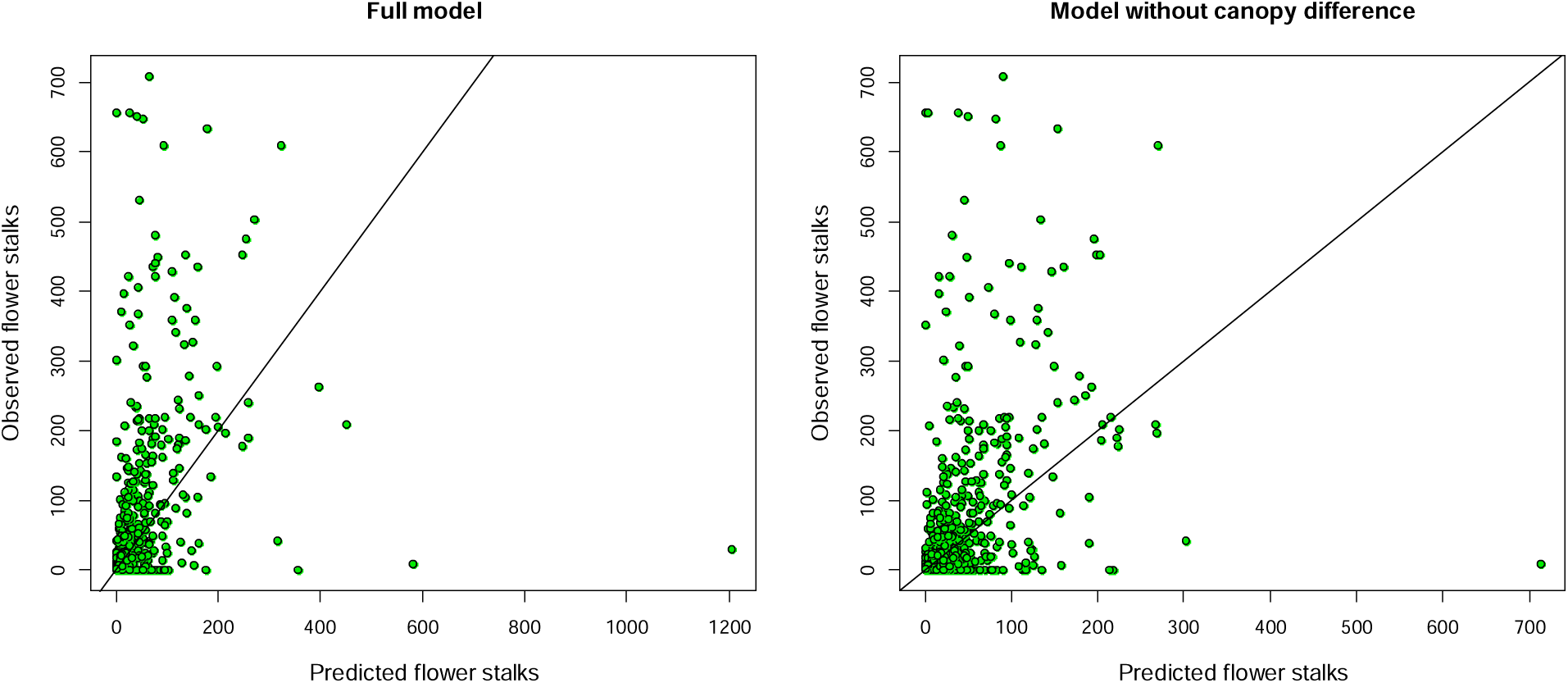
Model performance compared between the full model and the model without seasonal differences in canopy height between June and September. Predicted values were calculated out-of-sample using four-fold cross-validation.

### Prediction across sites

The model predicting flower stalk production performed better when evaluated using a random 4-fold cross-validation scheme compared to a spatially explicit leave-one-site-out approach. Under random 4-fold cross-validation, the model achieved a mean absolute error (MAE) of 100, with a 90% credibility interval (CI) ranging from 89 to 126. In contrast, the leave-one-site-out evaluation, which tests the model’s ability to generalize to new spatial contexts, resulted in a higher MAE of 125 and a much wider 90% CI (91 to 889), indicating substantial variation in predictive performance across sites. These results suggest that while the model performs consistently when trained and tested on spatially intermixed data, its predictive accuracy and reliability decrease when extrapolated to entirely new sites.

### Prediction across years

When evaluated using a random 3-fold cross-validation scheme, the model predicting flower stalk production achieved a median mean absolute error (MAE) of 108, with a 90% credibility interval (CI) ranging from 93 to 157 flower stalks. In contrast, performance declined substantially under a year-left-out cross-validation approach, which tests the model’s ability to generalize across time. In this temporally explicit scheme, the median MAE increased to 354 flower stalks, with an extremely wide 90% CI (125 to 10,590 flower stalks), indicating high variability and poor generalization to new years that lack training data. Compared to the earlier spatial validation results—where the model’s MAE increased moderately from a median of 100 (random 4-fold) to 125 (site-left-out)—the temporal validation shows a more dramatic loss of accuracy and consistency. This result indicates that, at least for our predictive model, temporal extrapolation poses a greater challenge for generalization than spatial extrapolation.

## Discussion

Identifying reproductive potential of wild plants at landscape scales will support both ecological research and land management needs. Our goal was to develop a low-cost workflow for predicting individual sagebrush flower stalks with UAV imagery. We found that an off-the-shelf RGB UAV could predict flower stalk production, with mean absolute error ∼100 flower stalks, well below the maximum number produced by large plants (>700 stalks). Structural variables, derived from structure-from-motion, were key to the success of our approach. Model performance improved with hurdle models that explicitly accounted for the high proportion of plants with zero flower stalks in our dataset. The cost-effectiveness of mapping sagebrush flower stalk production using UAV imagery depends on how well existing models can be transferred to new sites and years with minimal field data. We found mixed evidence for model transferability: predictive accuracy declined moderately when models were applied to sites that differed in topography and sagebrush genetic diversity, but dropped substantially when models were applied to different years. These findings highlight both the promise of using low-cost UAV imagery to estimate plant reproductive effort at landscape scales and the limitations of applying models without site- or year-specific training data.

Our results add to a growing body of evidence that high-resolution remotely sensed data can be applied to estimate flower production in wild plant populations (Smigaj and Gaulton 2021, Lee et al. 2023, Torresani et al. 2023). In contrast to previous studies, our work took place over an elevational gradient that included significant intraspecific variation, including multiple ploidy levels and different subspecies (Grossfurthner et al. 2023). The demonstrated ability to predict flower stalk production for genetically-diverse sagebrush plants is promising for scaling up our methodology to many other study sites across the Great Basin, where there is an urgent need to identify plants within locally adapted genotypes that are resilient to climate change and collect seed from them (Baughman et al. 2019).

Our first step in predicting flower stalk production, which, as expected included many zeroes, was to develop models that could account for high variability in our count data. We tested three models, a negative binomial generalized linear model, a zero-inflated model, and a hurdle model. The negative binomial model accounts for overdispersion but not excess zeroes. In contrast, the zero-inflated and hurdle models accommodate more zeroes than expected under a standard negative binomial distribution (Feng 2021). The hurdle model also allowed zero occurrence to be predicted using covariates. Both zero-inflated and hurdle models outperformed the negative binomial GLM, highlighting the importance of explicitly modeling reproductive failure. The hurdle model provided the best fit, indicating that reproductive failure could be predicted by model covariates derived from UAV imagery. This model reflects the biological reality that zero reproductive output may result from both ecological constraints (e.g., environmental stressors) and stochastic processes (Hassett and McGee 2017). Our results underscore the value of considering data-generating processes, including differentiating reproductive failure from counts of reproductive success, when developing predictive models for plant demography.

Structure-from-motion (SfM) was essential to our workflow, from identifying individual plants to model predictions. The canopy height model derived from SfM enabled us to segment individual shrubs across the landscape and match spectral and structural characteristics of shrub canopies to field data. For plants that successfully reproduced, structural variables, including edge-to-area ratio, mean canopy height, and seasonal differences in canopy height, had stronger effects than spectral variables. This finding is consistent with a previous study showing that lidar-derived crown volume was more predictive of fruit-tree yield than spectral variables (Chen et al. 2022). In sagebrush steppe ecosystems, low-cost SfM offers comparable accuracy to more expensive airborne lidar systems (Delparte, in prep). A caveat to our finding that structural variables outweighed spectral variables in our models is that we only included bands in the visible spectrum; multispectral or hyperspectral systems may have much better capacity to detect subtle changes in plant demography (Caughlin et al. 2016, Lee et al. 2023). Nonetheless, our results demonstrate that structure-from-motion can generate meaningful predictors of reproductive output, emphasizing the power of this technology for monitoring and modeling plant demography in drylands (Olsoy et al. 2024).

A structural variable that we expected to have a strong impact on predictive capacity for flower stalk production was seasonal differences in canopy height from June (the beginning of the growing season) to September (when flowers are produced). We found mixed support for the value of seasonal data on structure. In a model with all covariates, canopy height differences across season had strong effects on flower stalk production (*Figure. 5*). Nevertheless, out-of-sample predictions improved when variables related to canopy height differences were removed from the model. Our interpretation of these results is that UAV imagery collected across seasons is potentially a useful predictor of sagebrush flower stalk production. These results contrast with research from agricultural systems, where multi-temporal imagery typically improves predictions of crop yield (Yang et al. 2022, Camenzind and Yu 2024), emphasizing the increased challenge of accounting for interannual variation in natural systems. A worthwhile topic for further research would be to assess ideal collection dates for within-season image collection to maximize prediction. However, for immediate applicability of our methods, imagery from a single time point during peak flowering appears sufficient to predict the number of flower stalks on individual sagebrush plants.

Evaluating the transferability of predictive models built from UAV imagery is critical, as collecting field data for model training remains one of the most resource-intensive aspects of remote sensing applications (Amputu et al. 2024). When we tested spatial transferability using a leave-one-site-out approach, model performance declined moderately compared to random cross-validation, suggesting that spatial extrapolation is feasible if imperfect. This result was surprising, given the substantial differences in topography, soil type, and genetic composition of sagebrush plants between the four sites in our study area (Grossfurthner et al. 2023), and it points to the potential for scaling up our approach across heterogeneous rangelands. In contrast, model performance declined much more severely under year-left-out cross-validation. Because interannual variation strongly influences reproductive output, additional training data may be required when applying our model to novel years. An important caveat to the comparison between model transferability between sites and years is that sites were treated as random effects, allowing site-specific responses, whereas years were modeled as fixed effects due to limited sample size (n = 3). This difference in treatment likely contributed to the apparent gap in transferability across space versus time. More generally, our results are in line with a recent study that showed limited transferability of UAV-based predictions of dryland plant cover, richness, height and productivity, under different climate regimes (Blackburn et al. 2025). High-resolution UAV imagery may be particularly sensitive to interannual variation that alters relationships between ecological variables and remotely sensed measurements. Improving UAV model transferability remains a key challenge to avoid the need for extensive field data collection each time a drone is flown.

We anticipate that our results will have immediate applicability for monitoring population dynamics in sagebrush steppe. Integrating individual-level measurements across landscape-level gradients to develop population models is a long-standing goal in ecology (Gurevitch et al. 2016) that is directly relevant to understanding cross-scale variation in big sagebrush population dynamics (Germino et al. 2018, Zaiats et al. 2023). Our previous work has demonstrated the feasibility of UAV-based measurements of big sagebrush survival and growth (Olsoy et al. 2024) and seedling recruitment (Zaiats et al. 2024). This paper establishes the capacity to measure flower stalk production from UAVs, an advance that will enable full life cycle measurements of big sagebrush demography from aerial imagery. Landscape-level population models fit to UAV-based demographic rate measurements will enable spatially-explicit estimates of population viability, sensitivity to management interventions, and forecasts of population growth and decline. These advancements will enhance our ability to restore and conserve big sagebrush populations across the complex landscapes of the Great Basin.

### Limitations

Trade-offs between spatial scale, accuracy, and applicability are inherent to remote sensing (Jochems et al. 2021). We elected to fly UAVs at a relatively low altitude to capture fine-scale shrub structure. Flying at a higher altitude would enable users to sample a broader area, but would result in lower resolution. Even with our high-resolution imagery, including high overlap between photos to ensure detailed point clouds, we found that nearly half of field points could not be matched to canopy segments. This result is due to the complex structure of high-density shrublands, potentially leading to split canopies and crowns in segmentation products. Our solution to these errors was to select model parameters for segmentation that more accurately detect large shrubs, at the cost of missing smaller shrubs (Silva et al. 2016). For our application, identifying shrubs with highest fecundity, the loss of smaller shrubs from the dataset is unlikely to impact our results. However, for other use cases, optimizing segmentation algorithms to balance tradeoffs between shrub counts and accurate estimates of plant size may be necessary (Olsoy et al. 2024).

A major application for our methods will be UAV-based identification of highly fecund sagebrush plants for seed collection. However, we note that flower stalk production may not necessarily result in viable seed production, which depends on pollen reception and maturation of seeds, among other processes (Jacquemyn et al. 2010). Furthermore, selecting plants for seed collection based on measures of fecundity alone risks selecting traits that tradeoff with stress tolerance and other adaptations, potentially undermining long-term success of restoration projects (Pizza et al. 2021). To avoid these pitfalls, a useful future study would be to relate flower stalk count to number of sagebrush seeds produced and other demographic rates. Spatial maps of seed production across sagebrush crowns could then be combined with existing seed dispersal kernels for sagebrush plants (Applestein et al. 2022), enabling highly accurate maps of potential natural regeneration.

### Implications

Conservation and restoration of plant populations in rangeland ecosystems increasingly depends on tools that can inform adaptive management under changing environmental conditions. Our study shows that UAV imagery can estimate reproductive effort in a foundational shrub species with reasonable accuracy, even across individuals that differ widely in ploidy and subspecies identity. The framework we present offers rangeland managers a scalable method to assess shrub reproductive potential, with potential applications in planning restoration treatments, tracking vegetation recovery, and identifying resilient genotypes. Future work linking flower stalk production to viable seed yield and long-term plant performance will further improve the utility of UAV-based monitoring for sagebrush conservation. As UAV and open-source processing technologies continue to improve, their integration into rangeland monitoring programs will enhance our capacity to detect demographic patterns, guide seed sourcing strategies, and ultimately support sustainable management of sagebrush ecosystems.

## Acknowledgements

This publication was made possible by the NASA FINESST Program #80NSSC21K1638 and the National Science Foundation under award numbers OIA-1757324 and BIO-2207158. We thank the staff at Castle Rocks State Park for facilitating access to our study sites.

