## Appendix S1 for "Drone-Based Monitoring of Reproductive Potential in a Foundational Shrub Species"

**Appendix S1: Additional details on drone imagery collection and processing**

For article titled: **Drone-Based Monitoring of Reproductive Potential in a Foundational Shrub Species**

Ryan Scott Wickersham^1^, Megan E. Cattau^1^, Jennifer S. Forbey^1^, Valorie Marie^1^, Andrii Zaiats^1^, Donna M Delparte^2^, T. Trevor Caughlin^1^*

^1^ Boise State University, Boise, ID, USA

^2^ Idaho State University, Pocatello, ID, USA

*RGB image collection*

We collected RGB imagery using a DJI Mavic 2 Pro equipped with a 20MP Hasselblad RGB sensor. FAA licensed pilots conducted all flights and ensured compliance with all state, federal, and local regulations. Flights occurred between 10:00 and 14:00 (solar noon) during June and September of 2021-2023. Flights were planned to occur on days with clear or uniformly overcast conditions to minimize shadow effects on shrub canopies, which are critical for generating accurate structure-from-motion (SfM) reconstructions.

To ensure year-to-year consistency, mission plans were built from KML polygons in the WGS84 UTM Zone 12N projection, derived using QGIS. We applied a 10–20 m buffer to each plot to improve image overlap and quality. These polygons were uploaded to an iPhone via Map Pilot Pro, where cross-grid flight plans with an additional diagonal transect were created to capture images from multiple angles. Flights were parameterized to achieve a ground sampling distance of 1 cm (Table A), with forward and side overlap set to 70% and 85%, respectively. The camera was tilted 5° off nadir to improve 3D reconstruction of canopy structures.

To ensure year-to-year consistency, mission plans were built from KML polygons in the WGS84 UTM Zone 12N projection. We applied a 10–20 m buffer to each polygon to improve image overlap and quality. We used Map Pilot Pro to create cross-grid flight plans with an additional diagonal pass to capture photos at different angles, with the goal of improving SfM products. Flights were parameterized to achieve a ground sampling distance of 1 cm (Table A), with forward and side overlap set to 70% and 85%, respectively. The camera was tilted 5° off nadir to improve 3D reconstruction of canopy structures. Terrain awareness was used using default DEMs from Map Pilot Pro, allowing flights to keep the same above ground altitude without making any significant changes in overlap of images. We exported flight plan polygons to an iPhone for use in the field.

Ground control points (GCPs) were placed prior to each flight, targeting areas of variable slope and elevation. At least five GCPs were deployed per plot, typically four near the corners and one central point, all within 10 m of plot boundaries. GCP coordinates were collected using a Topcon RTK GPS unit for high spatial accuracy during post-processing. Imagery was stored on-board the UAV’s SD card. Flight metadata included records of wind conditions, cloud cover, and any mission deviations.

*Table A. Key flight parameters flown in June and September from 2021-2022 using Map Pilot Pro.*

| **Sensor** | **Above Ground Level (AGL)** | **Ground sampling distance** | **Camera Angle** | **Crossgrid** | **Flight speed** | **Forwardlap/Sidelap** |
| --- | --- | --- | --- | --- | --- | --- |
| 20MP Hasselblad | ~41 meters | 1.0cm | 5+ off NADIR (90 degrees) | Yes | 2m/s | ~70/85 |

*ODM RGB SfM processing*

Image processing was conducted using OpenDroneMap (ODM), an open-source photogrammetry platform that generates orthomosaics, digital surface models (DSMs), digital elevation models (DEMs), and 3D point clouds. Since ODM requires JPEG format, original DNG images were batch-converted via Windows PowerShell, preserving metadata for georeferencing. GCP coordinates were corrected to WGS84 and formatted in a CSV file with fields for GCP label, latitude, longitude, and elevation (decimal degrees). Using the GCP Editor Pro module in ODM, JPEG images and GCP data were imported. Markers were manually placed on each GCP within the imagery to ensure alignment. Once verified in WebODM, processing began with outputs projected in the WGS84 UTM Zone 12N coordinate system. Processing settings for the Castle Rocks sites are detailed in Table B.

*Table B. Parameters used for processing all orthomosaics from imagery collected at Castle Rocks State Park.*

Options: auto-boundary:true, camera-lens:brown, dem-resolution:1, depthmap-

resolution:2400, dsm:true, feature-quality:ultra, mesh-octree-depth:12, mesh-size:400000,

min-num-features:80000, optimize-disk-space:true, orthophoto-resolution:1, pc-

quality:high, pc-sample:0.0005, pc-tile:true, skip-3dmodel:true

*Canopy height model*

We constructed CHMs by subtracting the digital terrain model (DTM) from the DSM, derived from WebODM point clouds. Processing was done in CloudCompare using a cloth simulation filter (CSF), which differentiates ground and above-ground points by inverting the point cloud and interpolating terrain beneath shrub canopies (Zhang et al. 2016). Terrain classification relied on CSF parameters and additional Boolean logic to define ground versus non-ground returns. Outlier filtering and point-to-ground distance calculations refined the point cloud for CHM generation. Final CHMs were exported as CSVs containing X, Y, and Z coordinates for each 1 cm² pixel.

*Individual crown segmentation*

Segmentation of individual shrub crowns was performed using the lidR package in R. CHMs were divided into 10-meter tiles, and shrub apexes were detected using the lmf function and a geometric function of the form:

 F(x) = a·tan(x·b) + c·x

We used a geometric function of the form F(x) = a·tan(x·b) + c·x to detect shrub tops. The parameters in this equation were adjusted for each site to account for differences in shrub density and morphology. The parameter a controlled the degree of clumping, with higher values reducing the likelihood of shrub crowns merging. The parameter b adjusted the sensitivity of detection to neighboring shrubs, influencing how closely individual plants could be identified near larger neighbors. Lastly, c defined the search radius and was optimized to improve detection of larger shrubs within each site.

After detecting canopy tops, crowns were segmented using the silva2016 algorithm (Silva et al. 2016), which considers shrub width-to-height ratios and spatial exclusion rules. Exclusion thresholds were minimized to ensure complete capture of large crowns, as these are most likely to support high flower stalk densities. Parameters were standardized across sites for consistent crown delineation.
